# Neurochemical imaging reveals changes in dopamine dynamics with photoperiod in a seasonally social vole species

**DOI:** 10.1101/2025.05.11.652581

**Authors:** Jaewan Mun, Kelley C. Power, Annaliese K. Beery, Markita P. Landry

## Abstract

Studying dopamine signaling in non-model organisms is crucial for understanding the broad range of behaviors not represented in traditional model systems. However, exploring new species is often hindered by a scarcity of tools suitable for non-genetic models. In this work, we introduce near-infrared catecholamine nanosensors (nIRCats) to investigate dopamine dynamics in meadow voles, a rodent species that exhibits distinct changes in social behavior and neurobiology across photoperiods. We observe increased dopamine release and release site density in social voles under short photoperiods, suggesting adaptations linked to environmental changes. Moreover, pharmacological and extracellular manipulations demonstrate that social voles exhibit heightened responsiveness to dopamine-increasing interventions and resilience against suppressive conditions. These findings highlight a significant association between dopamine signaling and photoperiod-driven changes in social behavior and establish nIRCats as an effective tool for expanding our understanding of dopamine dynamics across non-model organisms.

## Introduction

Significant breakthroughs in neuroscience have emerged from the study of diverse species from the barn owl to the sea slug, reinforcing August Krogh’s famous declaration: “for many problems there is an animal on which it can be most conveniently studied” (1, 2). Studies of diverse organisms also enable comparisons across species, improving our understanding of shared mechanisms and of species-specific adaptations (3–6). Nowhere is this more evident than in the study of social behavior, which can be highly variable between even closely related species (7–9). Although the study of diverse species is crucial, it is accompanied by methodological challenges, as many tools and techniques available in model organisms are genetically based and challenging to apply outside of the model organisms for which they have been developed. The development of non-genetically encoded neurochemical sensors that can be applied in wide-ranging species promises to advance research on behaviors exhibited only by ‘weird’, non-traditional species. Here we validate and apply synthetic catecholamine sensors to investigate photoperiod-driven changes in dopamine release, reuptake, and diffusion in the striatum of meadow voles (*Microtus pennsylvanicus*), a non-model organism that offers a rare opportunity to study pathways underlying seasonal changes in social dynamics. Using our tool, we discover previously uncharacterized dopamine signaling differences in voles under short photoperiods in which they exhibit increased social interactions, indicating a potential role for changing dopamine in mediating this shift.

Dopamine, a neuromodulatory catecholamine most often associated with reward and pleasure, also plays important roles in regulating social behavior. Striatal dopamine in particular is elevated after mating, facilitates social bonding following mating in some species, and is involved in social reward processing in a number of species, including humans (10, 11). However, dopamine release and modulatory dynamics at the synaptic level that may support alterations in social behaviors remain largely unknown. Many current methods used to examine dopamine dynamics are limited by spatial and/or temporal resolution. Microdialysis, although capable of measuring extracellular dopamine with high specificity, has poor spatial and temporal resolution. Fast-scan cyclic voltammetry, while able to quantify extracellular dopamine at high temporal resolution, is not capable of visualizing dopamine in high spatial resolution. Genetically encoded dopamine-based sensors have overcome many of these challenges and are capable of imaging at high temporal resolution but are often limited to use in model organisms such as mice and rats and have yet to achieve spatial resolutions enabling single release site quantification. We recently developed near-infrared catecholamine nanosensors (nIRCats) that are capable of imaging dopamine with high spatiotemporal resolution (12, 13). nIRCats enable direct visualization of dopamine volume transmission by labeling the brain’s extracellular space with release site resolution, and because of their synthetic nature, they can be readily used in diverse, non-model organisms.

Meadow voles provide a unique opportunity to study mechanisms supporting social behavior because they exhibit predictable transitions in grouping behavior within a single species in response to environmental cues (14). Meadow voles are seasonally social: females are solitary and territorial in summer months but come together in mixed-sex groups in the winter (15, 16). In the wild, photoperiod is the primary cue many mammals use to determine the season (17, 18) and is sufficient in lab settings to induce behavioral and physiological phenotypes, such as sociability, associated with each season. A parallel transition is triggered by photoperiod in the lab (Fig. 1A): in long photoperiods (‘solitary photoperiods’; 14h light : 10h dark) female meadow voles interact less with and are less affiliative toward conspecifics than their counterparts in short photoperiods (‘social photoperiods’; 10h light : 14h dark) (Fig. 1B,C) (19, 20). Many neurobiological pathways change in concert with the transition between photoperiods and social phenotypes, including changes in multiple neuropeptide signaling pathways (21–23). In prairie voles, a closely related monogamous vole species, dopamine signaling is integral to the formation and maintenance of social bonds with mate partners (24–28). Photoperiodic changes in dopamine signaling have been documented in mice and chipmunks (29–31), and may contribute to changes in behavior. In meadow voles, however, seasonal/photoperiodic changes in dopamine signaling have never been examined, nor have release-site differences in dopamine signaling in any species

**Figure 1.**
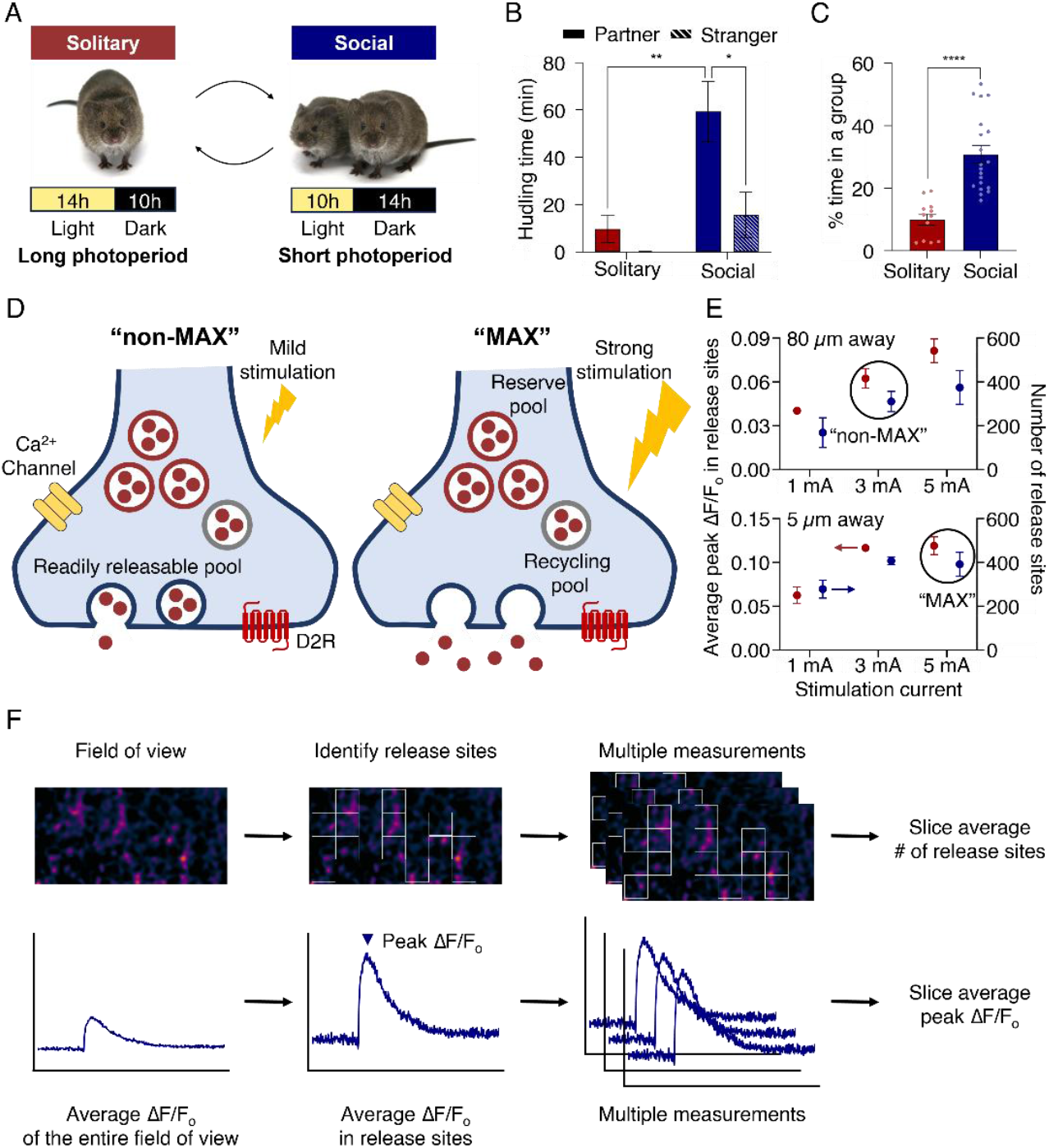
Schematic of Experimental Design. (A) Meadow voles in long, summer-like photoperiods (‘solitary photoperiod’) are less social than those in short, winter-like photoperiods (‘social photoperiods’) in the lab. (B) Long photoperiod (red) voles spend less time huddling with a cagemate peer or an unknown stranger than short photoperiod (blue) voles during a partner preference test. Data redrawn from Beery et al., 2008. (C) Long photoperiod (red) voles spend less time in groups of three or four during one week in a multi-chambered apparatus than short photoperiod (blue) voles. Data from Power et al., 2025. (D) Schematic of dopaminergic neurons under mild (left, “non-MAX”) and strong (right, “MAX”) electrical stimulation. More dopamine is released from the readily releasable dopamine pool with stronger electrical stimulation. (E) Change in average peak ΔF/F0 (red, left y-axis) and the number of release sites (blue, right y-axis) as a function of current, with the stimulator positioned at 80 μm (top) and 5 μm (bottom) from the field of view. (F) Schematic of data analysis methods used to examine the average number of release sites and the average peak ΔF/F0 per release site in a brain slice. The number of release sites and the peak ΔF/F0 correlate with the putative number of dopamine-releasing synaptic sites and the amount of dopamine released from each site, respectively. (n.s. not significant, *p<0.05, **p<0.01, ***p<0.005, ****p<0.001)

In the present study, we validated the use of nIRCats in meadow voles to investigate whether striatal dopamine release dynamics differed between voles with social and solitary behavioral phenotypes. We found that short photoperiod (social phenotype) voles exhibited higher dopamine release than long photoperiod voles, as reflected by both a greater number of dopamine release sites and a larger amount of dopamine released per site. We also investigated the effects of sex and regional variability within the striatum on dopamine dynamics. Finally, we modulated dopaminergic signaling using sulpiride (D2 receptor antagonist), quinpirole (D2 receptor agonist), and extracellular Ca^2+^ to explore how voles with different social phenotypes respond differently to these key regulators of dopamine signaling. In short photoperiod voles, dopamine release was more strongly enhanced in response to dopamine-increasing manipulations. In contrast, in long photoperiod voles, dopamine release was more strongly suppressed in response to dopamine-decreasing manipulations. These data provide evidence for increased striatal dopamine in social versus solitary phenotypes and establish nIRCats as a valuable tool to measure dopamine dynamics at the synaptic level in non-model organisms.

## Results

Voles in long photoperiods exhibit summer-like solitary phenotypes, whereas those housed in short photoperiods exhibit winter-like social phenotypes (Figure 1A-C). We sought to understand how dopamine signaling might be affected by photoperiodic changes in meadow voles, as day length is the primary cue regulating a major transition in their social behavior. To investigate how photoperiod impacts dopamine signaling, we synthesized dopamine-sensitive probes, nIRCats, following established protocols, and used these probes to label the extracellular space of brain slices prepared from adult voles exposed to the above-mentioned photoperiods with custom near-infrared microscopy(12, 13). We hypothesized that nIRCats would enable us to image changes in dopamine signaling through the brain extracellular space induced by different photoperiod exposures. We further hypothesized that there may be subtle changes in dopamine signaling, depending on whether neuronal synapses are stimulated with a mild stimulus to release predominately their readily releasable pool of dopamine, or whether synapses are promoted to release intracellular dopamine reserves with a strong stimulus to release most intracellular dopamine.

To study these phenomena, we first imaged dopamine release prompted by a single-pulse electrical stimulation applied under two different stimulation strength conditions to induce varying levels of dopamine release by adjusting the stimulation current and the distance between the imaging field of view and the electrode (Figure 1D). When a stimulation current greater than 0.3 mA was applied with the electrode positioned close (5 µm) to the field of view, the response of nIRCats became saturated and did not change with further increases in current beyond 0.3 mA (Figure 1E). This saturated response may be due to the release of all available readily-releasable dopamine vesicles from overstimulation (Figure 1D). We referred to this condition as ‘MAX,’ as it represents the maximum observable dopamine release in a field of view upon stimulation. In contrast, when the electrode was positioned 80 µm away from the imaging field of view, changes in stimulation current significantly affected the amount of dopamine release (Figure 1E). In this ‘non-MAX’ condition, only a portion of vesicles are released from the readily-releasable dopamine pool (Figure 1D). We chose 0.3 mA with the stimulator placed 80 µm away from the field of view as a representative ‘non-MAX’ condition to investigate non-saturated dopamine release.

One key advantage of this approach is the ability to image dopamine release and reuptake with high spatiotemporal resolution. Labeled brain slices were analyzed using custom Python code (32), which identified brain regions where dopamine fluorescence changes exceeded twice the standard deviation of baseline fluctuations upon stimulation. These dopamine “hotspots” were designated as regions of interest (ROIs), which we term dopamine release sites. Recent studies identified such release sites as tyrosine hydroxylase-positive axonal varicosities co-localized with the presynaptic protein Bassoon (33, 34). Thus, the number of release sites in our labeled slices may closely correspond to the number of single synaptic dopamine release sites. The degree of fluorescence modulation in each release site was quantified as ΔF/F_o_, defined as the fluorescence change (F-F_o_) divided by baseline fluorescence (F_o_), where peak ΔF/F_o_ corresponds to the amount of dopamine released from each release site. Therefore, the total amount of released dopamine is a function of both the number of release sites and the peak ΔF/F_o_ in each release site, and we used these two parameters to characterize dopamine release. Multiple measurements were performed for each brain slice to calculate the slice-averaged number of release sites and peak ΔF/F_o_ (Figure 1F).

We focused on differences between female meadow voles housed in short or long photoperiods as females have a more pronounced shift in social behavior with season and photoperiod than males. Since both females and males in short photoperiods huddle in mixed sex groups, we also investigated the effect of sex within short photoperiods for both ‘MAX’ and ‘Non-MAX’ conditions. Because grouping and photoperiod shift in tandem under natural conditions, it is helpful to mirror this natural combination in the lab, as well as to isolate the effect of photoperiod under uniform housing conditions. For both MAX and Non-MAX conditions we ran cohorts with naturalistic housing: short photoperiod voles were housed in groups of four and long photoperiod voles were housed alone. In the MAX stimulation group, we included additional cohorts of pair-housed voles in short and long photoperiods to assess the impact of day length alone.

### Max Stimulation Effects

We first applied ‘MAX’ stimulation to the dorsomedial striatum (DMS) of short photoperiod and long photoperiod voles across housing conditions to compare the size of the releasable dopamine pool between solitary and social phenotypes. Housing had no detectable influence on any of the outcomes measured compared to photoperiod (Figure S1, Supplementary Materials), thus data were pooled into short- and long-photoperiod groups for subsequent analyses. ‘MAX’ stimulation evoked an instantaneous increase in nIRCat fluorescence in both female long (peak ΔF/F_o_ = 0.105 ± 0.006, n = 15 brain slices) and short photoperiod voles (peak ΔF/F_o_ = 0.155 ± 0.012, n = 13 brain slices), as shown in Figure 2A-C. This fluorescence response was reversible, with fluorescence levels returning to baseline after 15 seconds through a combination of dopamine reuptake and diffusion away from the release site (Figure 2A,B). The peak ΔF/F_o_ was then calculated for each release site, rather than the entire field of view, to better quantify the relative amount of dopamine released as the release site density across experimental groups. Interestingly, short photoperiod voles exhibited significantly larger peak ΔF/F_o_ than long photoperiod voles, indicating that more dopamine was electrically evoked from dopamine release sites of social voles. Next, we compared the number of release sites between long photoperiod and short photoperiod voles (Figure 2D). Similar to the ΔF/F0 magnitude, short photoperiod voles had a larger number of release sites than long photoperiod voles (number of release sites of short photoperiod voles = 452 ± 9, n = 13 brain slices, number of release sites of long photoperiod voles = 392 ± 20, n = 15 brain slices), suggesting that short photoperiod voles possess a higher release site density, with each release site releasing more dopamine (ΔF/F_o_) than long photoperiod voles.

**Figure 2.**
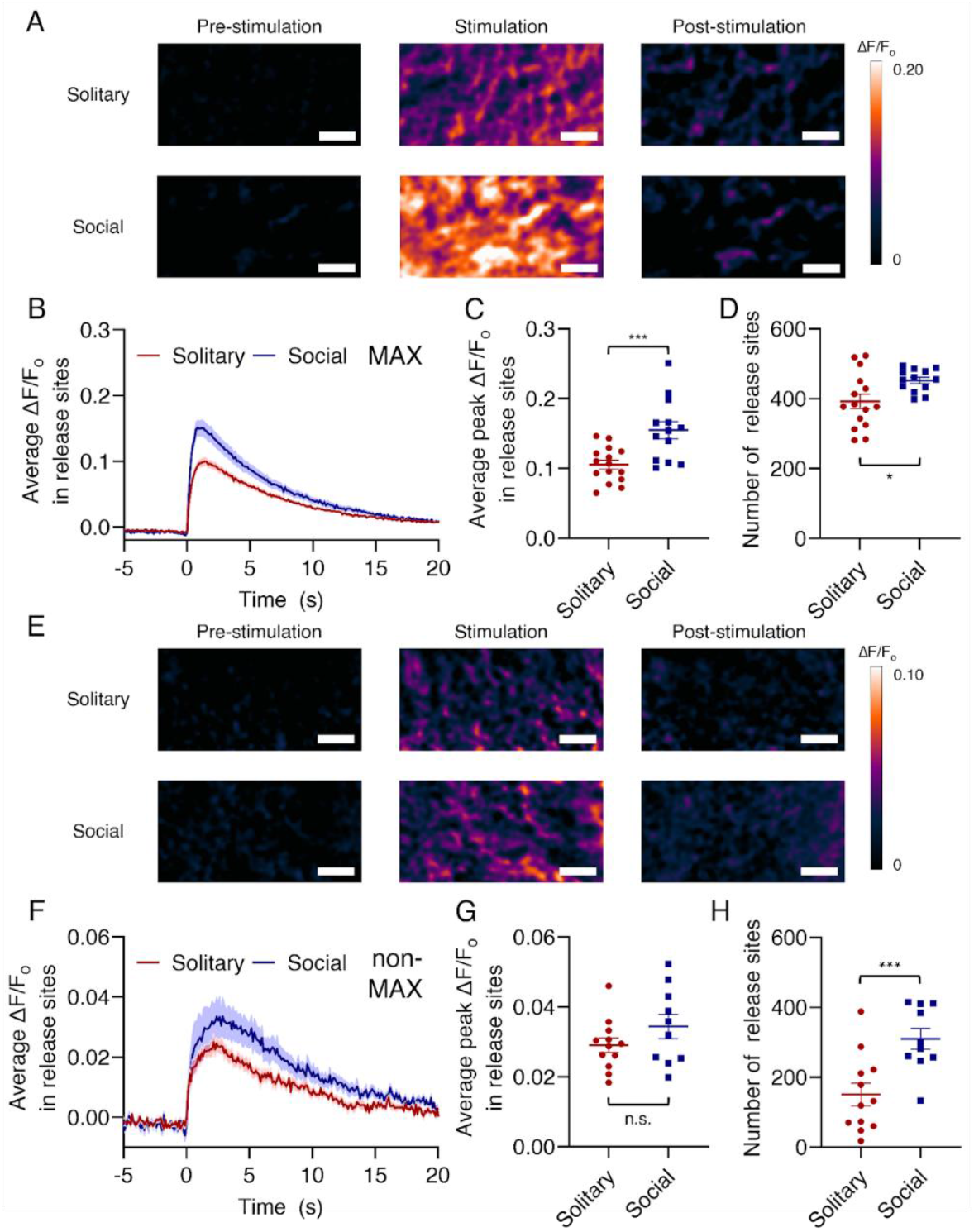
Comparison Across Social and Solitary Photoperiods. In slice, imaging electrically-evoked dopamine in the DMS of long photoperiod (red) and short photoperiod (blue) voles following a 0.5 mA electrical stimulation applied adjacent to the field of view (MAX, a-d) and a 0.3 mA electrical stimulation applied 80 μm away from the field of view (non-MAX, e-h). (A) ΔF/F0 response images within the same field of view, (B) average ΔF/F_o_ in release sites time trace (solid line) with the standard error (shadow), (C) average peak ΔF/F0 in each release site, and (D) number of release sites in the MAX condition. (E) ΔF/F_o_ response images within the same field of view, (F) average ΔF/F_o_ in release sites time trace (solid line) with the standard error (shadow) (G) average peak ΔF/F_o_ in each release site, and (H) number of release sites in the non-MAX condition. “Pre-stimulation”, “Stimulation”, and “Post-stimulation” represent nanosensor fluorescence before, immediately following, and after electrical stimulation, respectively.” (Scale bar: 10 µm, n.s.: not significant, *p<0.05, **p<0.01, ***p<0.005)

### Non-MAX Stimulation Effects

We next applied ‘Non-MAX’ stimulation to the DMS of a new cohort of both short and long photoperiod voles. ‘Non-MAX’ stimulation resulted in an increase in dopamine release in in both short and long photoperiods, with much less dopamine release observed in both cohorts compared to the ‘MAX’ stimulation condition, as shown in Figure 2E-G. Under non-MAX stimulation conditions, nIRCats again showed clear reversibility, with their fluorescence returning to baseline as dopamine is reuptaken and diffuses away from the release site within 10 seconds of stimulation.

Under these non-MAX stimulation conditions, short photoperiod voles showed a trend toward increased dopamine release per site compared to long photoperiod voles, although the difference was not statistically significant. Social voles showed a peak ΔF/F_o_ of 0.034 ± 0.003 (n = 10 brain slices), while long photoperiod voles exhibited a peak ΔF/F_o_ of 0.029 ± 0.002 (n = 12 brain slices) (Figure 2G). Combined with the observations from those of MAX stimulation results, we concluded that dopamine release sites in social voles contain a larger amount of dopamine, as shown in Figure 2C, but mild electrical stimulation evoked a similar amount of dopamine release from each site of both short and long photoperiods, as shown in Figure 2G. Under non-MAX stimulation conditions, short photoperiod voles again showed a significantly larger number of release sites (310 ± 29, n = 10 brain slices) than long photoperiod voles (150 ± 31, n = 12 brain slices) (Figure 2H). When compared to ‘MAX’ stimulation, short photoperiod voles exhibited a 31.4% reduction in the number of dopamine release sites, while long photoperiod voles showed a 61.7% reduction under the ‘non-MAX’ condition. Therefore, we propose that short photoperiod voles not only have more dopamine release sites available, as shown in Figure 2D, but their release sites are also more readily activated by electrical stimulation than those of long photoperiod voles, as shown in Figure 2H. Taken together, our data show that under mild stimulation conditions, short photoperiod voles exhibited more activated dopamine release sites than long photoperiod voles, though release sites effluxed similar quantities of dopamine per site across both cohorts.

Lastly, it is known that voles of both sexes housed in a short, social photoperiod form selective preferences for known (vs. unknown) same-sex conspecifics and huddle in groups (9, 35). To identify whether sex differences in dopamine release exist, we assessed male and female meadow voles housed in the short photoperiod. Because of their similar social behavior in this photoperiod, we hypothesized that males and females would not show significant differences in dopamine release. Upon applying ‘MAX’ and ‘non-MAX’ electrical stimulation in the DMS, indeed neither ΔF/F_o_ nor the number of release sites differed significantly between short photoperiod male and female voles (Figure S2,3, Supplementary Materials).

### Regional Variability within the Dorsal Striatum

Having observed differences in dopamine signaling as a function of stimulation strength, we next studied whether there were subregional differences in dopamine signaling within the dorsal striatum of short photoperiod voles. The dorsal striatum can be divided into two regions-the dorsomedial striatum (DMS) and the dorsolateral striatum (DLS). While both subregions are implicated in motor control, they serve distinct behavioral functions. Dopamine in the DMS plays a critical role in goal-directed behaviors and behavioral flexibility, while dopamine in the DLS is responsible for habitual behaviors and navigation (36, 37). To this end, dopamine was electrically evoked in these two regions within the same brain slices under ‘MAX’ condition. Under ‘MAX’ stimulation, no significant difference in peak ΔF/F_o_ in release sites was observed between the DMS and the DLS, although the two regions clearly differed in the number of release sites. (Figure S4, Supplementary Materials) Our results showed that dopaminergic neurons in the DMS contain a larger number of readily releasable dopamine vesicles than those in the DLS, but the maximum amount of the dopamine that can be released from each vesicle is similar in both striatal sub- regions.

### Effects of D2 autoreceptor manipulation

Having examined differences in dopamine release dynamics across sexes, social behaviors, and striatal subregions, we next investigated how dopamine D2 autoreceptor manipulation affects evoked dopamine release in short and long photoperiods. D2 autoreceptors play a crucial role in regulating dopamine signaling by suppressing dopamine release and synthesis when necessary, maintaining homeostasis in the brain’s dopamine system. These receptors are primarily located on the presynaptic terminals of dopaminergic neurons (Figure 1D). When dopamine is released into the synaptic cleft, it can bind to these autoreceptors, which then send a feedback signal to the neuron to reduce further dopamine release and synthesis. Due to their critical function, D2 autoreceptors have been widely studied in the context of dopamine-related psychiatric diseases such as Parkinson’s disease and schizophrenia. (38–40)

It has previously been demonstrated that nIRCats are compatible with dopamine pharmacology, including the use of sulpiride (a D2 autoreceptor antagonist) and quinpirole (a D2 autoreceptor agonist) (12). Based on our above data showing differences in dopamine release site density and amount as a function of photoperiod, we hypothesized that long photoperiod and short photoperiod voles may have differing levels of dopamine receptor sensitivity to pharmacological intervention. We first examined the effects of sulpiride, a D2 autoreceptor antagonist, on electrically evoked dopamine release in long photoperiod and short photoperiod voles. After imaging dopamine release in ACSF, 5 µM of sulpiride was bath-applied for 30 minutes and electrically evoked dopamine release was then imaged in the same field of view. For this experiment, ‘non-MAX’ stimulation was used exclusively, as overstimulation could override any effects of the applied pharmacological agents.

We found that a 5 µM concentration of sulpiride significantly increased dopamine release in the DMS, as shown in Figure 3A,B. ΔF/F0 per release site in long photoperiod voles increased by 74%, from a peak ΔF/F_o_ of 0.031 ± 0.002 pre-sulpiride to 0.054 ± 0.006 post-sulpiride (n = 5 brain slices), while in short photoperiod voles, the peak ΔF/F_o_ increased by 92%, from 0.037 ± 0.005 pre-sulpiride to 0.072 ± 0.012 post-sulpiride (n = 5 brain slices) (Figure 3C). These results confirm that antagonism of presynaptic D2 autoreceptors using sulpiride effectively increases the amount of dopamine released from each dopamine release site. Furthermore, these results suggest that social phenotype voles exhibit a higher sensitivity to dopamine autoreceptor antagonism relative to their solitary counterparts.

**Figure 3.**
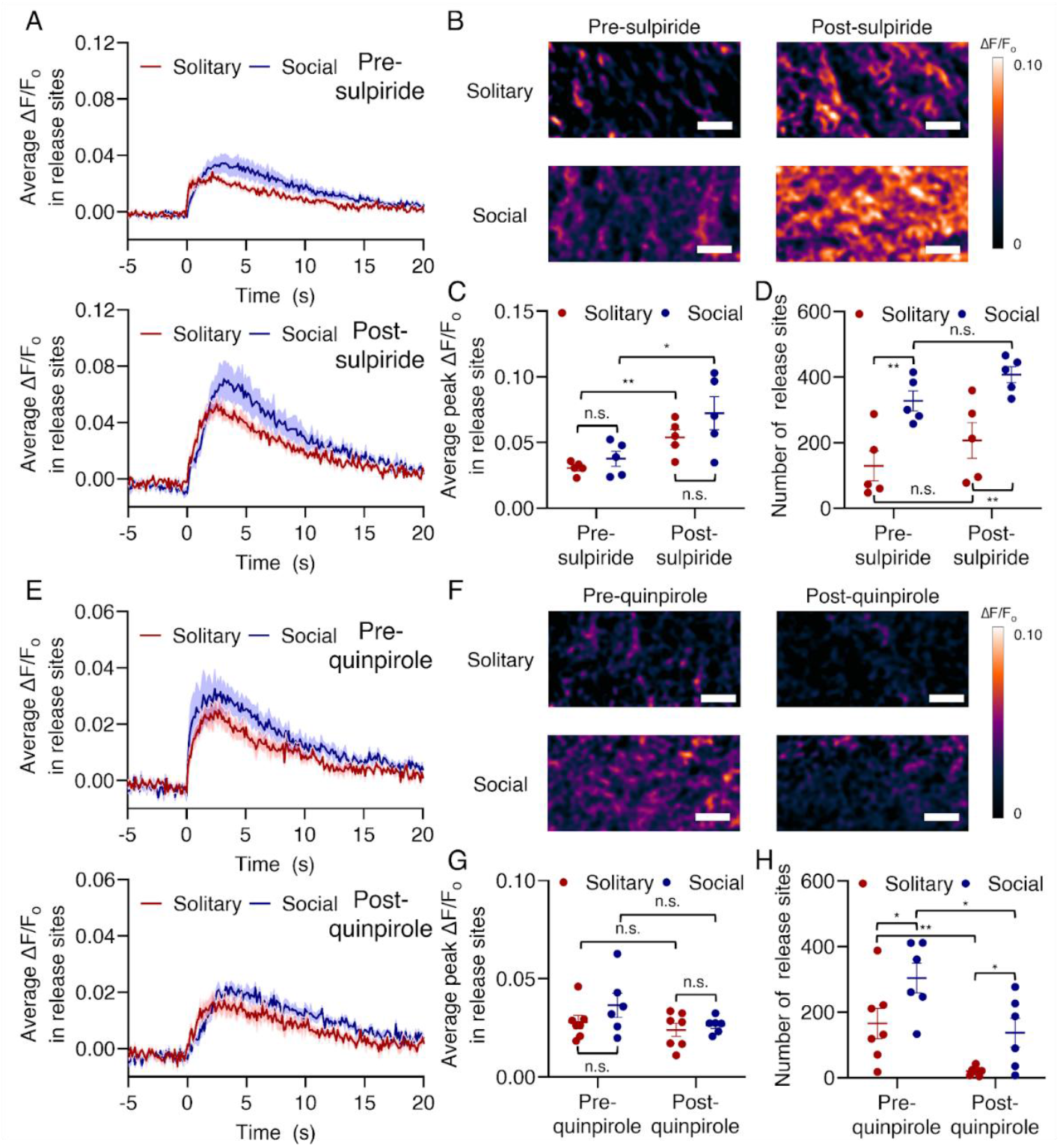
Effects of D2 autoreceptor manipulation. In slice, effects of D2-autoreceptor antagonist (sulpiride, A-D) and agonist (quinpirole, E-H) on electrically-evoked dopamine release of solitary (long photoperiod; red) and social (short photoperiod; blue) voles. (A) average ΔF/F_o_ in each release site’s time trace (solid line) with the standard error (shadow), (B) ΔF/F_o_ response images, (C) average peak ΔF/F_o_ and (D) number of release site following a 0.3 mA single-pulse electrical stimulation in the DMS of short and long photoperiod voles before and after 5 µM sulpiride treatment. (E) average ΔF/F_o_ in each release site’s time trace (solid line) with the standard error (shadow), (F) ΔF/F_o_ response images, (G) average peak ΔF/F_o_ and (H) number of release sites following a 0.3 mA single-pulse electrical stimulation in the DMS of short and long photoperiod voles before and after 5 µM quinpirole treatment. (Scale bar: 10 µm, n.s.: not significant, *p<0.05, **p<0.01, ***p<0.005)

While the application of 5 µM sulpiride clearly increased ΔF/F_o_, or the amount of dopamine released per release site in both long and short photoperiod voles, its effects were less evident in the number of release sites (Figure 3D). In long photoperiod voles, the number of release sites increased by 60%, from 129 ± 46 pre-sulpiride to 207 ± 54 post-sulpiride (n = 5 brain slices), whereas in short photoperiods, the release site number rose by 24%, from 327 ± 30 pre-sulpiride to 407 ± 24 post-sulpiride (n = 5 brain slices). Interestingly, after 5 µM sulpiride treatment, most release sites in short photoperiod voles were activated, with the number of release sites approaching that seen during ‘MAX’ stimulation conditions (452 ± 9, n = 13 brain slices) (Figure 2D). However, even with 5 µM sulpiride, most dopamine release sites in long photoperiod voles remained deactivated, and the number of release sites fell short compared to those measured with ‘MAX’ stimulation conditions (392 ± 20, n = 15 brain slices). These results suggest that dopamine autoreceptors in social phenotype voles are more responsive to sulpiride treatment, reaching near- maximal activation of release sites, whereas dopamine release in solitary phenotype voles remains less responsive, with many sites still inactive. This difference implies that social vole dopaminergic neurons may develop a greater capacity for dopamine release upon stimulation than their solitary phenotype peers.

Next, we investigated how quinpirole, a D2 autoreceptor agonist, affects electrically evoked dopamine release in the DMS of long photoperiod and short photoperiod voles. Dopamine release upon ‘non-MAX’ electrical stimulation was measured in the same field of view before and after bath application of 5 µM quinpirole. Surprisingly, although quinpirole substantially reduced integrated fluorescence across the field of view, the amount of dopamine released from each release site decreased only marginally in both long photoperiod and short photoperiod voles (Figure 3E-G). The ΔF/F_o_ measured from release sites changed from 0.028 ± 0.004 and 0.036 ± 0.007 pre- quinpirole to 0.024 ± 0.004 and 0.026 ± 0.002 post-quinpirole for long (n = 7 brain slices) and short (n = 6 brain slices) photoperiods, respectively. Instead, the major source of the clear decrease in dopamine response was the reduction in the number of dopamine release sites. The number of release sites in short photoperiod voles decreased by 55%, from 307 ± 48 pre-quinpirole to 137 ± 49 post-quinpirole (n = 6 brain slices), while the release site number in long photoperiod voles was reduced by 88%, from 165 ± 50 pre-quinpirole to 20 ± 5 post-quinpirole (n = 7 brain slices). Although short photoperiod voles retained nearly half of their release sites after quinpirole application, dopamine release sites in long photoperiod voles were mostly deactivated, suggesting that solitary voles may have a higher sensitivity to D2 autoreceptor agonism relative to their social counterparts, potentially due to elevated D2 autoreceptor expression or heightened functional responsiveness. This increased sensitivity could indicate an inhibitory mechanism that constrains dopamine release more stringently in solitary voles, aligning with reduced dopaminergic signaling.

Based on our measurements, we identified several characteristics of how D2 autoreceptors are differentially manipulated by dopamine receptor pharmacology, hinting at autoreceptor expression or sensitivity differences in social versus solitary phenotype voles. Although both drugs affect the number of dopamine release sites and the amount of dopamine released from each site to some extent, they operate through different mechanisms. Specifically, sulpiride is more effective at increasing the amount of dopamine released from each site, while quinpirole is better at deactivating release sites. In addition, we found that social phenotype voles respond more sensitively to sulpiride, which increases dopamine, but are less responsive to quinpirole, which decreases dopamine.

### Extracellular Ca^2+^ Effects

Finally, we investigated how extracellular Ca^2^+ affects electrically-evoked dopamine release in long photoperiod and short photoperiod voles. The release of neurotransmitters such as dopamine from presynaptic dopaminergic neurons is highly dependent on the influx of Ca^2^+ through voltage-gated calcium channels (Figure 1D). Consequently, the amount of dopamine released is influenced by the concentration of extracellular Ca^2^+. We hypothesized that long photoperiod and short photoperiod voles might differentially respond to varying concentrations of extracellular Ca^2^+, as suggested by our prior data showing differing sensitivities of each photoperiod to MAX vs. non-MAX stimulation conditions.

For this experiment, short and long photoperiod vole acute brain slices were prepared and incubated in Ca^2^+-deficient ACSF buffer before imaging. This environment ensures that prior Ca^2^+ exposure does not influence dopamine release upon stimulation. Electrically-evoked dopamine was imaged under ‘non-MAX’ stimulation conditions in the Ca^2^+-deficient buffer. After this initial imaging, Ca^2^+ was added to final concentrations of 2 mM and 5 mM, and dopamine release was imaged in the same field of view. In 0 mM Ca^2^+ buffer, we observed that dopamine release was highly suppressed in both long photoperiod and short photoperiod voles, as expected (Figure 4). Long photoperiod voles exhibited a negligible amount of dopamine release, with a peak ΔF/F_o_ of 0.009 ± 0.004 and an average of 8 ± 3 release sites (n = 6 brain slices). Compared to dopamine release in regular ACSF buffer (Figure 2E-H), most dopamine release sites were deactivated (a 94% reduction in the number of release sites) in the absence of Ca^2^+, and the few active sites released significantly smaller amounts of dopamine relative to standard 2mM Ca^2^+ buffer conditions. Interestingly, short photoperiod voles still maintained some of their release sites even in a Ca^2^+-deficient environment, with a peak ΔF/F_o_ of 0.016 ± 0.007 and an average of 68 ± 30 release sites (n = 5 brain slices). These observations suggest that social voles exhibit higher resistance to Ca^2^+ deficiency in maintaining dopamine release, even in the absence of extracellular Ca^2^+.

**Figure 4.**
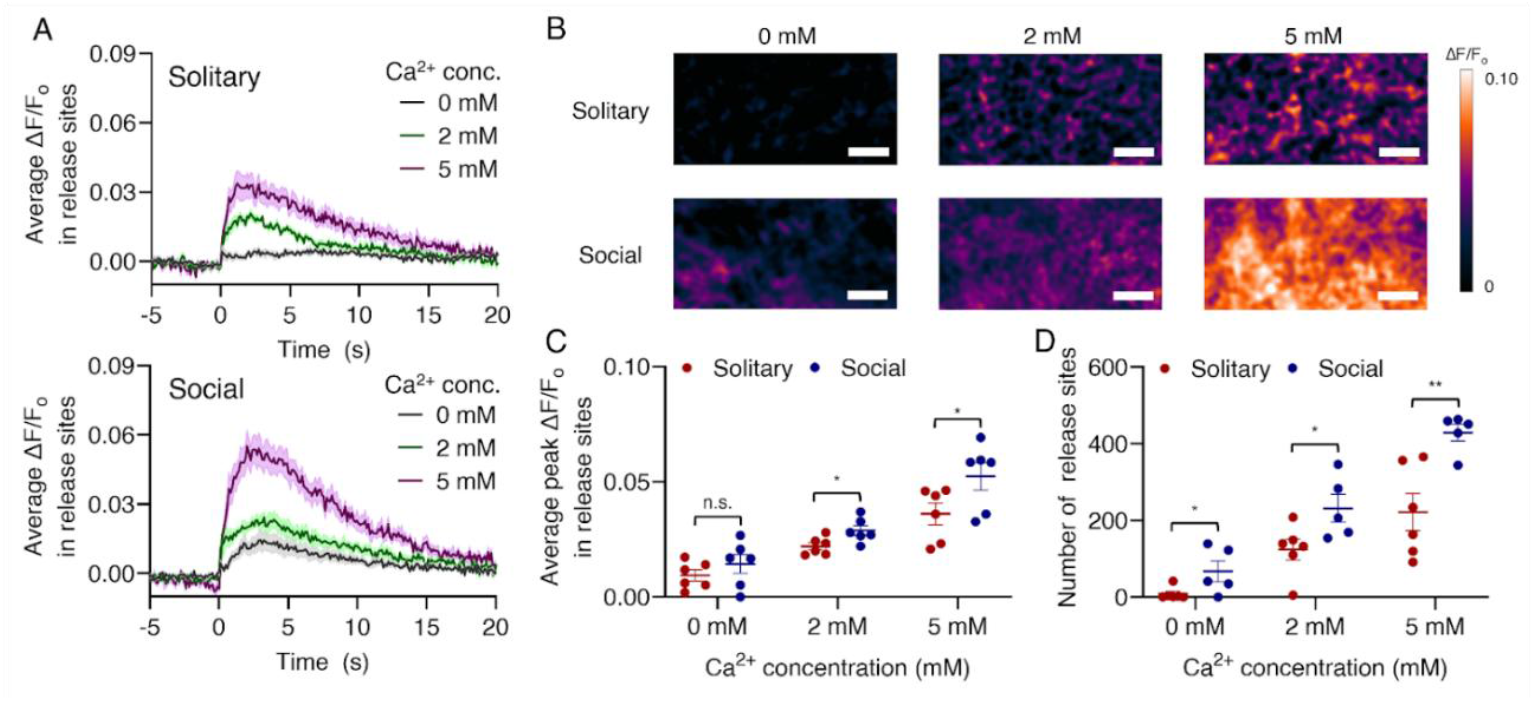
Effects of extracellular Ca^2+^. In slice, effects of extracellular Ca^2+^ on electrically-evoked dopamine release of long photoperiod (red) and short photoperiod (blue) voles. (A) average ΔF/F_o_ in each release site’s time trace (solid line) with the standard error (shadow), (B) ΔF/F0 response images, (C) average peak ΔF/F_o_ and (D) number of release sites following a 0.3 mA single-pulse electrical stimulation in the DMS of short and long photoperiod voles with various concentrations of Ca^2+^. (Scale bar: 10 µm, n.s.: not significant, *p<0.05, **p<0.01)

Next, a 2 mM Ca^2^+ buffer was supplied for 30 minutes, and upon stimulation, dopamine release became clearly observable, confirming that extracellular Ca^2^+ plays a critical role in dopamine release, as expected (Figure 4B). Interestingly, under these 2 mM Ca^2^+ extracellular calcium conditions, short photoperiod voles exhibited both a higher peak ΔF/F_o_ (0.029 ± 0.003 for social, n = 5 brain slices, and 0.022 ± 0.002 for solitary, n = 6 brain slices) and a greater number of release sites compared to long photoperiod voles (231 ± 36 for social, n = 5 brain slices, and 124 ± 27 for solitary, n = 6 brain slices) (Figure 4C,D). It should be noted that when imaging was performed in regular ACSF without prior exposure to a 0 mM Ca^2^+ environment, the difference in peak ΔF/F_o_ between solitary and social voles was insignificant (Figure 2G,H). These results indicate that social phenotype voles recover more quickly from Ca^2^+-deficient conditions than their solitary counterparts. However, neither solitary nor social phenotypes showed full recovery of dopamine release to the levels observed when measured without prior exposure to a 0 mM Ca^2^+ environment. Next, a 5 mM Ca^2^+ buffer was supplied for another 30 minutes, which further increased both the peak ΔF/F_o_ and the number of release sites upon electrical stimulation. Short photoperiod voles exhibited a peak ΔF/F_o_ of 0.056 ± 0.006 and 429 ± 22 release sites (n = 5 brain slices). Interestingly, 5 mM of extracellular Ca^2^+ fully activated available dopamine release sites, whereby the observed number of release sites was comparable to that measured under ‘MAX’ stimulation conditions (Figure 2D). Although at 5 mM Ca^2^+ long photoperiod voles also exhibited increases in peak ΔF/F_o_ (0.036 ± 0.005) and the number of release sites (222 ± 48, n = 6 brain slices) relative to lower Ca^2^+ conditions, the release site numbers remained well below values observed under ‘MAX’ stimulation (Figure 2D).

In summary, based on all observations from varying extracellular Ca^2^+ concentrations, we confirmed that Ca^2^+ plays a significant role in gating dopamine release, with increasing Ca^2^+ levels leading to higher peak ΔF/F_o_ and a greater number of release sites. Our findings are consistent with previous studies showing that elevated extracellular calcium concentrations can increase the likelihood of neurotransmitter vesicle release, expand the size of the readily releasable pool, and shift inactive boutons into more active states (41, 42). Similar to what we observed with D2 autoreceptor drugs, short photoperiod voles showed lower dopamine suppression under zero Ca^2^+ concentrations, which reduces dopamine release, and demonstrated faster enhancement in dopamine release as Ca^2^+ concentrations increased. Taken together, our Ca^2^+ data provides further evidence that social voles either synthesize higher levels of intracellular dopamine, and/or develop a higher synaptic vesicle count that is more readily releasable than their solitary counterparts.

## Discussion

Non-traditional species enable the study of multiple behaviors not found in model organisms. For example, meadow voles undergo a unique transition in social behavior in response to photoperiod that allows us to examine how changes in the brain contribute to changes in social behavior. Studying non-model organisms comes with a trade-off, however: many of the tools available in model organisms are genetically based and may not work across non-model species. In this study, we use nIRCats to explore dopamine dynamics in meadow voles, a unique species that exhibits changes in social or solitary phenotypes depending on photoperiod. Using photoperiod-induced transitions in vole social behavior, we observed striking differences in synaptic-scale striatal dopamine signaling between social and solitary vole phenotypes.

Across a wide variety of experimental conditions including stimulation profiles, Ca^2+^ concentrations, agonist and antagonist applications, evoked dopamine was altered in a consistent, photoperiod-specific manner. Voles housed in short, social photoperiods had both a greater number of dopamine release sites and higher dopamine release than voles housed in long, solitary photoperiods in the MAX condition. Under mild (‘non-MAX’) stimulation, short photoperiod voles still activated more release sites than long photoperiod voles, but there was no difference in the amount of dopamine each site released. Additionally, we observed that social voles demonstrated heightened resistance to conditions that reduce dopamine release, such as quinpirole treatment or lack of extracellular calcium. Winter groups are essential for survival and thermoregulation for meadow voles (15, 43, 44). It is possible that group living may be supported not only by elevated dopamine but also resistance to reduced dopamine, ensuring voles remain in a group throughout the winter. To the same end, social voles showed greater sensitivity to agents that increase dopamine release, such as sulpiride and 5 mM extracellular calcium, further supporting the role of dopamine in regulating social behaviors and the possibility of dopamine synthesis upregulation or differences in autoreceptor sensitivity induced by photoperiod.

While social behavior changes markedly across photoperiods, this change is part of a larger suite of physiological and behavioral traits that shift as animals prepare for the transition to a winter environment. In winter and short photoperiods, voles are reproductively quiescent, weigh less and decrease food intake (45), and are more active than those housed in long photoperiods (20). Increased dopamine levels in short photoperiods may support this increase in locomotion and may also contribute to changes in motivational states related to reproduction, feeding and other behavioral drives.

Interestingly, many rodent studies of the interactions between dopamine and photoperiod find evidence of *lower* dopamine in short photoperiods relative to long photoperiods (29–31), opposite the findings in this study. Humans, however, show increased striatal dopamine during the fall and winter short photoperiods (46), in line with findings in the present study, and there is some evidence in rodents that photoperiodic effects on dopamine vary between species and brain regions (31), further highlighting species-specific variation. Furthermore, social contexts can also impact dopamine. Historically, studies of social isolation on dopamine have shown an association between isolation and increased dopamine response (47–49). In meadow voles, isolation isn’t a stressor for voles in long, ‘solitary’ photoperiods as it is in many other rodent species (50, 51). We tested voles in both matched and naturalistic housing conditions and found no differences by housing conditions; it’s possible that testing short photoperiod voles in unnatural solitary housing conditions would lead to similar changes in dopamine response.

The nIRCats used in this study were used in slice preparation, which is most suitable for detecting stable alterations in release-site scale dopamine signaling, e.g., across different environmental conditions such as photoperiod. Our results herein compare differences in the number of release sites, and the amount of dopamine released per release site, where some conditions induce changes in one but not both metrics. These results highlight the importance of release site scale dopamine imaging to measure nuanced differences likely unobservable using bulk measurement approaches. As we look towards the future of non-genomically encoded sensor use, future studies will benefit from nIRCats or other sensors that are capable of imaging non-model organisms *in vivo* to measure context-dependent dopamine signaling. Additionally, since dopamine can act as a neuromodulator and alter the activity of other neuropeptides like oxytocin and cortisol that are known to vary seasonally, sensors that can measure multiple molecules simultaneously would be beneficial to further elucidate these relationships.

The present findings not only validate the use of nIRCats in voles but also highlight the importance of dopamine in modulating seasonal behavioral transitions in meadow voles. This work paves the way for future studies of dopamine signaling in non-traditional organisms and demonstrates the potential of nIRCats for advancing our understanding of neurotransmitter dynamics across species.

## Materials and Methods

### Animal Subjects

Meadow voles were bred locally at UC Berkeley in long photoperiods (14h light: 10h dark). Within one week of weaning, voles were separated into experimental housing and either remained in long photoperiods, or were moved to short photoperiods (10h light: 14h dark). For the initial MAX and Non-MAX conditions and regional variability experiments, female meadow voles were housed in same-sex, age-matched pairs in both photoperiods. For all other conditions, male and female meadow voles were housed in same-sex, age-matched groups of four in short photoperiods (social phenotype), and females were housed alone in long photoperiods (solitary phenotype). Voles were all adults at the time of the experiment. All animal procedures were approved by the University of California Berkeley Animal Care and Use Committee.

### Materials

HiPco SWNTs were purchased from NanoIntegris (batch #27-104, diameter: 0.8-1.2 nm, length: 400-700 nm). (GT)_6_ oligonucleotides purified with standard desalting were purchased from Integrated DNA Technologies. Quinpirole and sulpiride were purchased from Tocris Bioscience. All other agents were purchased from Millipore Sigma.

### Sensor Synthesis and in vitro Characterization

Near-infrared catecholamine sensors were synthesized using previously reported protocols. Briefly, SWNTs and (GT)_6_ (2 mg each) were bath-sonicated (Branson Ultrasonic 1800) for 3 minutes in 1 mL of 0.1 M NaCl, followed by probe tip-sonication for 10 minutes (Cole-Parmer Ultrasonic Processor, 3-mm diameter tip, 5W power) in an ice bath. To remove non-dispered SWNTs, the resulting suspension was centrifuged at 16,000g for 60 minutes (Eppendorf 5418), and the supernatant containing stable (GT)_6_-SWNT was collected and used. The absorbance of the (GT)_6_-SWNT was measured at 632 nm (NanoDrop One, Thermo Scientific) to calculate the SWNT concentration of the suspension. (extinction coefficient ε = 0.036 (mg/L)^-1^ cm^-1^) The (GT)_6_- SWNT suspension was diluted to 5 μg mL^-1^ and 200 μg mL^-1^ for in vitro characterization and *ex vivo* slice experiments, respectively.

For in vitro characterization, fluorescence measurements were performed using a 20X objective on an inverted Zeiss microscope (Axio Observer D1) with a SCT 320 spectrometer (Princeton Instruments) and a liquid nitrogen-cooled InGaAs linear array detector (PyLoN-IR, Princeton Instruments). Aliquots of 99 μL of (GT)_6_-SWNT were placed in each well of a 384-well plate. The fluorescence spectra of (GT)_6_-SWNT were obtained before and after the addition of 1 μL of dopamine using a custom-built spectrometer and microscope. A 721 nm laser (Opto Engine LLC) was used to excite the nanosensor suspensions. nIRCats exhibit a 24-fold increase in fluorescence upon exposure to dopamine, with high selectivity against various neurochemicals in vitro (52).

### Brain Slice Preparation and Sensor Labeling

Brain slices were prepared using established protocols (12, 13, 53). Briefly, meadow voles were anesthetized under isoflurane and then intraperitoneally injected with a combination of ketamine (10 mg mL^-1^) and xylazine (5 mg mL^-1^) in saline (1 mL). While anesthetized, transcardial perfusion was performed with ice-cold cutting buffer (119 mM NaCl, 26.2 mM NaHCO_3_, 2.5 mM KCl, 1 mM NaH_2_PO_4_, 3.5 mM MgCl_2_, 10 mM glucose, and 0 mM CaCl_2_), followed by rapid brain dissection over the same buffer. The brain was mounted onto a vibratome (Leica VT 1000S) and coronally sliced into 300 μm thick sections. Sections containing the dorsal striatum were incubated for 30 minutes at room temperature in oxygen-saturated ACSF buffer (119 mM NaCl, 26.2 mM NaHCO_3_, 2.5 mM KCl, 1 mM NaH_2_PO_4_, 1.3 mM MgCl_2_, 10 mM glucose, 2 mM CaCl_2_). The slices were transferred to a small volume incubation chamber (Scientific Systems Design Inc., AutoMate Scientific) containing 5 mL of oxygen-saturated ACSF. nIRCat nanosensors were added to the incubation chamber (final concentration of 200 µg mL^-1^), and slices were incubated for 15 minutes. Then, the slices were rinsed with ACSF buffer to remove excess non-localized sensors.

### Dopamine Imaging in Acute Brain Slices

Slice imaging was performed using established protocols and a custom-built upright epifluorescent microscope (Olympus, Sutter Instruments) mounted onto a motorized stage (12, 13). A 785 nm laser (Opto Engine LLC) was used to excite nanosensors, which was expanded to a final diameter of ∼1 cm using a Keplerian beam expander with two plano-convex lenses (f = 25 and 75 mm; AR coating B, Thorlabs) The beam was passed through a fluorescence filter cube [excitation: 800 nm shortpass (FESH0800), dichroic: 900 longpass (DMLP990R), and emission: 900 longpass (FELH0900); Thorlabs] to a 60× Apo objective (numerical aperture, 1.0; working distance, 2.8 mm; water dipping; high nIR transmission; Nikon CFI Apo 60XW nIR). Emission photons collected from the sample were passed through the filter cube, were focused onto a two-dimensional InGaAs array detector [500 to 600 nm: 40% quantum efficiency (QE); 1000 to 1500 nm: >85% QE; Ninox 640, Raptor Photonics], and were recorded using the Micro-Manager Open Source Microscopy Software.

A bi-polar stimulation electrode (Platinum/Iridium Standard Tip, MicroProbes for Life Science) was positioned in the dorsal striatum using a 4x objective lens. A total of 600 frames were captured in the nIR using a 60x objective lens at 8 frames per second, with electrical stimulation applied after 200 frames of baseline. For the saturated condition, the electrode was placed directly next to the field of view, and a single-pulse 0.5 mA stimulation was applied for 1 millisecond. For the non-saturated condition, the electrode was placed 80 μm away from the field of view, and a single-pulse 0.3 mA stimulation was applied for 1 millisecond. nIRCats demonstrated fluorescence modulation in response to evoked dopamine release with high spatial (μm) and temporal (ms) resolution.

For drug experiments, imaging was initially conducted in plain ACSF. Then, either quinpirole or sulpiride (5 µM) was added to the imaging chamber through ACSF perfusion. The brain slice was incubated with either drug for 30 minutes before imaging continued in the same field of view. For calcium experiments, initial incubation occurred in ACSF without calcium, then calcium was later applied in concentrations of 2 mM and 5 mM. Brain slices were exposed to new calcium concentrations for 30 minutes before imaging.

### Image Processing and Data Analysis

Image files were processed using a custom Python application (https://github.com/NicholasOuassil/NanoImgPro). A field of view was divided into 25 × 25 pixel grids, and those with fluorescence modulation more than two times the standard deviation of baseline fluctuations were identified as ROIs, which we term release sites. Recent studies identified such release sites as tyrosine hydroxylase-positive axonal varicosities co-localized with the presynaptic protein Bassoon (33, 34). Thus, the number of release sites may closely correspond to the number of single synaptic dopamine release sites. ΔF/F0 of each release site was calculated as (F-F_o_)/F_o_, where F_o_ is the average intensity of baseline fluorescence and F is the dynamic fluorescence intensity. ΔF/F_o_ was averaged over release sites to draw an average ΔF/F_o_ in release sites as a function of time. The maximum ΔF/F_o_ of each release site was identified and averaged to calculate the average peak ΔF/F0 in release sites. The total amount of released dopamine is a function of both the number of release sites and the peak ΔF/F_o_ in each release site, and we these two parameters to describe electrically-evoked dopamine release.

## Data and materials availability

All of our data are available in the main text or in the supporting information. A custom image processing Python application is available online. (https://github.com/NicholasOuassil/NanoImgPro).

## Acknowledgements

This material is based upon work supported by the National Science Foundation Graduate Research Fellowship Program under Grant No. DGE 2146752 (K.C.P.). Any opinions, findings, and conclusions or recommendations expressed in this material are those of the author(s) and do not necessarily reflect the views of the National Science Foundation.

We acknowledge support of a National Science Foundation CAREER Award 2239635 (A.B.), a National Institutes of Health grant R01MH132908 (A.B.), a Burroughs Wellcome Fund Career Award at the Scientific Interface (M.P.L.), a Dreyfus Foundation Award (M.P.L.), the Philomathia Foundation (M.P.L.), an NSF CAREER Award 2046159 (M.P.L.), McKnight Foundation Award (M.P.L.), a Simons Foundation Award (M.P.L.), a Moore Foundation Award (M.P.L.), a Heising-Simons Fellowship (M.P.L.), a Brain Foundation Award (M.P.L.), a Polymaths award from Schmidt Sciences, LLC (M.P.L.). M.P.L. is a Chan Zuckerberg Biohub investigator.

