## Supplementary figures for "Neurochemical imaging reveals changes in dopamine dynamics with photoperiod in a seasonally social vole species"

### This PDF file includes:

Figs. S1 to S4

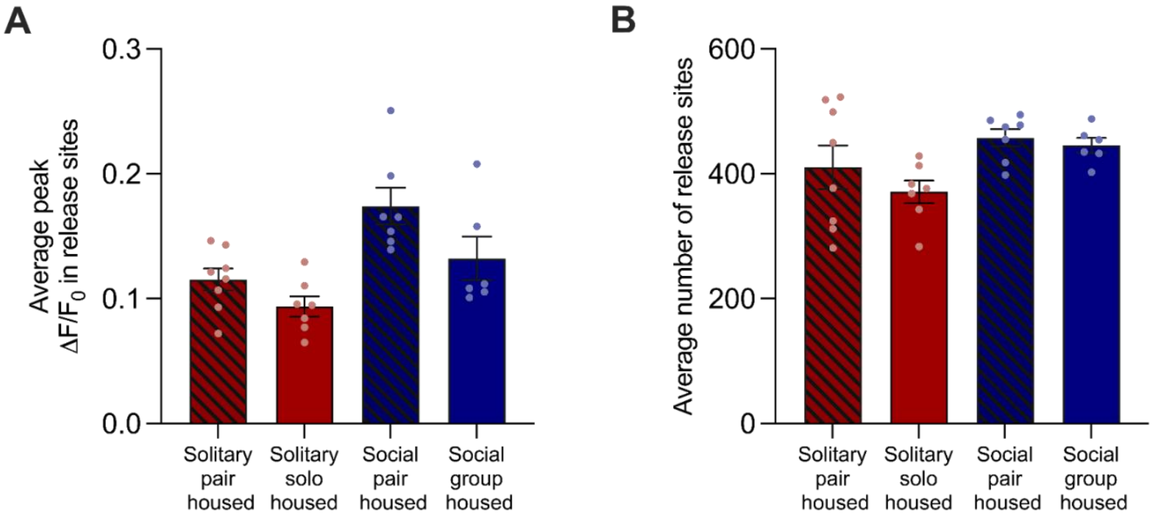


**Fig. S1. In slice, electrically-evoked dopamine in the DMS of female social (blue) and solitary (red) voles following a 0.5 mA electrical stimulation applied adjacent to the field of view.** (A) Average peak $\Delta F/F_0$ in dopamine release sites, and (B) number of dopamine release sites under 'MAX' stimulation conditions.

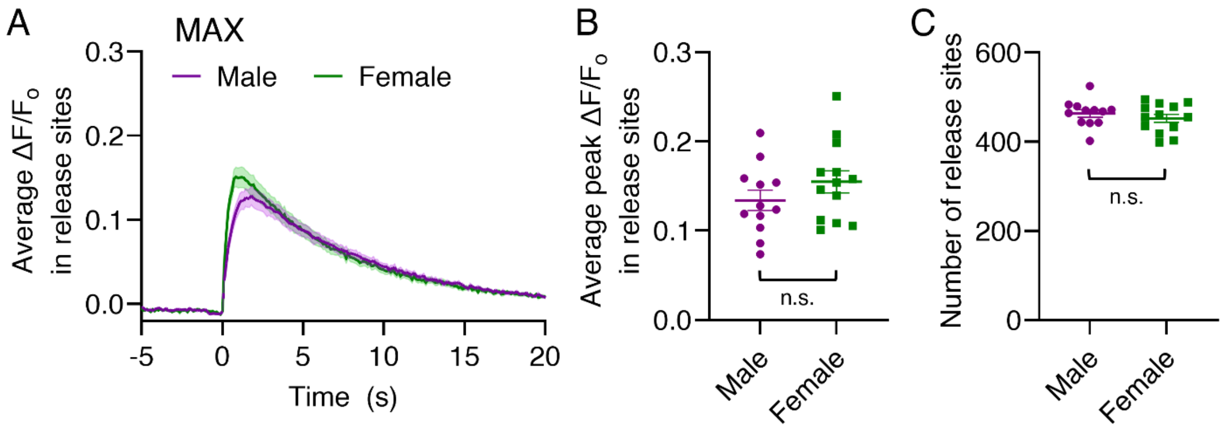


**Fig. S2. In slice, imaging electrically-evoked dopamine in the DMS of male (purple) and female social (green) voles following a 0.5 mA electrical stimulation applied adjacent to the field of view.** (A) average $\Delta F/F_0$ in release site time traces (solid line) with standard error (shadow), (B) average peak $\Delta F/F_0$ in release sites, and (C) number of release sites under 'MAX' stimulation conditions. (n.s. not significant)

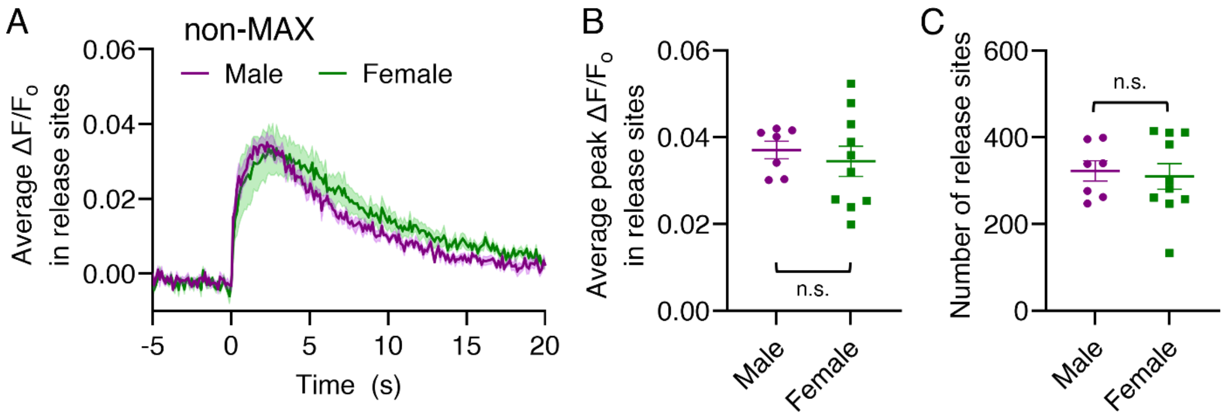


**Fig. S3. In slice, imaging electrically-evoked dopamine in the DMS of male (purple) and female social (green) voles following a 0.3 mA electrical stimulation applied 80 µm away from the field of view.** (A) average $\Delta F/F_0$ in release site time traces (solid line) with standard error (shadow), (B) average peak $\Delta F/F_0$ in release site, and (C) number of release sites under 'non-MAX' stimulation conditions. (n.s. not significant)

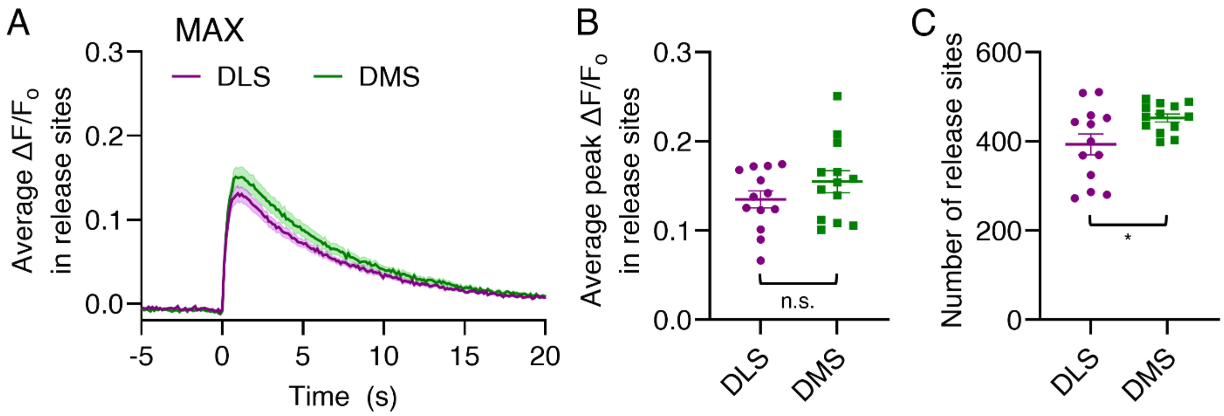


**Fig. S4. In slice, imaging electrically-evoked dopamine in the DLS (purple) and the DMS (green) of female social voles following a 0.5 mA electrical stimulation applied adjacent to the field of view.** (A) average $\Delta F/F_0$ in release site time traces (solid line) with standard error (shadow), (B) average peak $\Delta F/F_0$ in release sites, and (C) number of release sites under 'MAX' stimulation conditions. (n.s. not significant, *p<0.05)
